# Prevalence and Dynamics of Genome Rearrangements in Bacteria and Archaea

**DOI:** 10.1101/2024.10.04.616710

**Authors:** Carolina A. Martinez-Gutierrez, Louis-Marie Bobay

**Affiliations:** Department of Biological Sciences, North Carolina State University; Department of Earth Science, University of California Santa Barbara

**Keywords:** genome rearrangement, synteny, transposable elements, genome rearrangement rates, origin of replication

## Abstract

The genetic material of bacteria and archaea is organized into various structures and set-ups, attesting that genome architecture is dynamic in these organisms. However, strong selective pressures are also acting to preserve genome organization, and it remains unclear how frequently genomes experience rearrangements and what mechanisms lead to these processes. Here, we assessed the dynamics and the drivers of genomic rearrangements across 121 microbial species. We show that synteny is highly conserved within most species, although several species present exceptionally flexible genomic layouts. Our results show a rather variable pace at which genomic rearrangements occur across bacteria and archaea, pointing to different selective constraints driving the accumulation of genomic changes across species. Importantly, we found that not only inversions but also translocations are highly enriched near the origin of replication (*Ori*), which suggests that many rearrangements may confer an adaptive advantage to the cell through the relocation of genes that benefit from gene dosage effects. Finally, our results support the view that mobile genetic elements—in particular transposable elements—are the main drivers of genomic translocations and inversions. Overall, our study shows that microbial species present largely stable genomic layouts and identifies key patterns and drivers of genome rearrangements in prokaryotes.

**Significance statement:** Bacterial and archaeal genomes display stable architectures which ensures the preservation of fundamental cellular processes. However, large genomic rearrangements occasionally occur. Although most of these events are thought to be highly deleterious, they have the potential to lead to adaptive events. Here, we examined the general trends of the dynamic of prokaryotic genomes by exploring the occurrence of genome rearrangements across a broad diversity of bacterial and archaeal species. We find that genomes remain highly syntenic in most species over short evolutionary timescales, although some species appear particularly dynamic. Rearrangements are strongly biased, and most gene blocks are relocated near the origin of replication. We also measured remarkably variables rates at which genome rearrangements occur across species, and transposons and other mobile genetic elements appear to be the main drivers of these variations. Overall, this study provides a comprehensive picture of the dynamic of genome architecture across many microbial species.

## INTRODUCTION

Bacterial genomes are highly organized entities that evolve under the selection pressures imposed by functional constraints such as replication, transcription, DNA repair, and cell division (1). Over long timescales, genomes have evolved into diverse chromosomal set-ups (e.g., circular/linear chromosomes, single/multiple chromosomes) through processes mediated by homologous recombination, horizontal gene transfer, and mobile genetic elements (MGEs) (1–3). Multiple studies have shown that genomes are not simply the set of genes of an organism (3–5). Instead, genomes display a complex architecture whose organization is shaped and constrained by natural selection to facilitate cell processes (1, 3). For instance, genes involved in the same function are often organized into operons, and essential genes that are highly expressed tend to be found on the leading strand and in close proximity to the origin of replication (*Ori*) (3, 6). Genes located near *Ori* are highly expressed because this DNA region is present in two or more copies when the chromosome is actively replicating (i.e., gene dosage effect) (3, 7, 8). In addition, microbial genomes harbor polarized short DNA motifs that play essential roles for chromosome stability and duplication (3). For instance, *Chi* sites are eight-nucleotide-long motifs present on the leading strand of Enterobacteria, where they play a key role in DNA repair (9). In *Escherichia coli*, polarized motifs such as *KOPS* are found near the terminus of replication (*Ter*), and are essential for the proper segregation of the two chromosomes during the termination of DNA replication (3, 10, 11). In *E. coli*, the chromosome is structured into four macrodomains and two unstructured regions (12). This architecture is important for chromosome dynamics during the cell cycle and the shuffling of macrodomains and the motifs they contain is expected to deleteriously affect cell fitness (12, 13).

Due to the essential role of genome organization in bacteria, genomic rearrangements are thought to be rare because they often alter the architecture and polarity of the genome (3). For instance, genome inversions are expected to deleteriously affect cell fitness by reversing the polarized DNA motifs along the chromosome(s) (13). Other studies have suggested that—despite the constraints of chromosome architecture— certain types of rearrangements can be common in bacteria. It has been shown that *E. coli, Haemophilus influenzae*, and *Helicobacter pylori* undergo rearrangements due to recombination, but inversions tend to be symmetrical relative to the replication axis, which preserves strand polarity (14). However, studies focusing on the impact of genomic rearrangements on cellular fitness have led to contrasting conclusions. Several works have shown that genomic inversions can play a positive role in the ability of microbes to evade host defenses (4, 15, 16), but others suggest that these events lead to lower growth rates and even cell death (4, 17).

Current knowledge on genomic rearrangements is based on a narrow set of microbial clades. It remains unclear how frequently these events occur, what processes drive their occurrence, and to what extent they can impose deleterious, neutral, or beneficial selective pressures. In this study, we explore the prevalence and the potential impacts and drivers of genome rearrangements across a broad diversity of microbial species. By focusing our analyses on a large set of fully assembled genomes, we show that genomic rearrangements remain rare in the majority of the species analyzed, although their rates vary greatly across species. We found evidence that selection purges most of the rearrangements occurring in microbial genomes. Most rearrangements occurred near and symmetrically to *Ori*, suggesting that these events are less deleterious—or potentially beneficial—when occurring in this region. Interestingly, rearrangements near *Ori* are not restrained to inversions, but translocations are also enriched in this region. These results suggest that rearrangements are not predominantly composed of inversions occurring near *Ori*, which are thought to be less detrimental to the cell by preserving strand polarity. Rather, these patterns support the view that many rearrangements may be adaptive through the relocation of genes near *Ori* to benefit from increased expression through gene dosage effect. Finally, we present strong evidence showing that transposable elements and other MGEs are likely the main drivers of genome rearrangements in bacterial and archaeal species.

## RESULTS

### Identification of genomic rearrangements in bacterial and archaeal species

In order to explore the prevalence of genome rearrangements across a broad diversity of prokaryotic species, we collected high-quality genomes (100% complete and <5% contaminated) belonging to archaea and bacteria from the Genome Taxonomy Database v207 (GTDB; GTDB dataset) (18). Our GTDB dataset consisted of 52,543 genomes belonging to 5,077 and 142 bacterial and archaeal species, respectively (Supplementary File 1). Given the expected role of assembly level on genomic rearrangement identification, we obtained the assembly information for each genome from the National Center for Biotechnology Information. We selected the genomes that were fully assembled for further analyses, and species with fewer than five genomes were excluded. Our final dataset (Final dataset; Supplementary File 2) was composed of 5,238 genomes belonging to 118 bacterial species and three archaeal species.

For each species, we built the set of core genes (i.e., orthologous genes that are shared amongst most genomes of the species) using *CoreCruncher* (19), which is robust to the presence of paralogs and xenologs (see Methods). We then identified blocks of adjacent genes by comparing the order of core genes between genome pairs of the same species. We defined gene blocks as sets of consecutive and uninterrupted core genes between each genome pair, and only gene blocks with at least two core genes were considered to avoid biases due to the potential misassignment of gene orthology (see Methods). For each species, we selected one reference genome which was defined as the genome that led to the lowest average number of blocks in the species. The number of gene blocks between genome pairs was then used to estimate the number of genome rearrangements that have occurred within each species.

To avoid overestimates in the number of gene blocks due to incorrect assemblies, we assessed the impact of sequencing technology on gene block identification. Genomes generated through short-read sequencing (e.g., Illumina) can contain assembly errors as they are more complex to assemble relative to genomes generated from long-read sequencing technologies (e.g., PacBio and Oxford NanoPore). However, a statistical comparison showed that both sequencing technologies lead to similar estimates (Wilcoxon test, *P*=0.44, SI Appendix, Fig. S1). This finding indicates that short-read sequencing technologies do not appear to introduce more assembly errors than long-read sequencing technologies. Therefore, we used genomes generated by both, short- and long-read sequencing technologies for further analyses.

Overall, most of the bacterial and archaeal genomes analyzed harbored few gene blocks, indicating that the reorganization of the genome structure is infrequent within species (Fig. 1A). We found a median of three gene blocks per genome, with some genomes exceptionally harboring up to 132 blocks (Fig. 1A). A large fraction of the genomes in our dataset (32% of the total analyzed; Supplementary File 3) had only one gene block per genome, which is suggestive of the complete absence of genomic rearrangements in these genomes (Fig. 1A; Supplementary File 3). On average, the rearranged blocks in our dataset were composed of 184 genes (including accessory and core genes) and few blocks exhibited more than 1,500 genes (Fig. 1B). At the species level, most taxa had an average number of blocks below five (75% of the species considered; Supplementary File 4), with several species having an average of up to 43 blocks (Fig. 1C). We found seven species that did not exhibit any genomic rearrangements given that a single block of genes was inferred across all their genomes. These species were *Buchnera aphidicola, Treponema pallidum, Morganella morganii, Brevibacterium aurantiacum, Borreliela garinii, Borreliela burgdorferi, and Bordetella avium*. At the other end of the spectrum, *Aeromonas salmonicida, Bordetella pertussis, Treponema phagedenis, and Xanthomonas oryzae* had the highest number of gene blocks (>20 blocks) (Fig. 1C; Supplementary File 4), which reflects a higher occurrence of rearrangements in these species.

**Figure 1.**
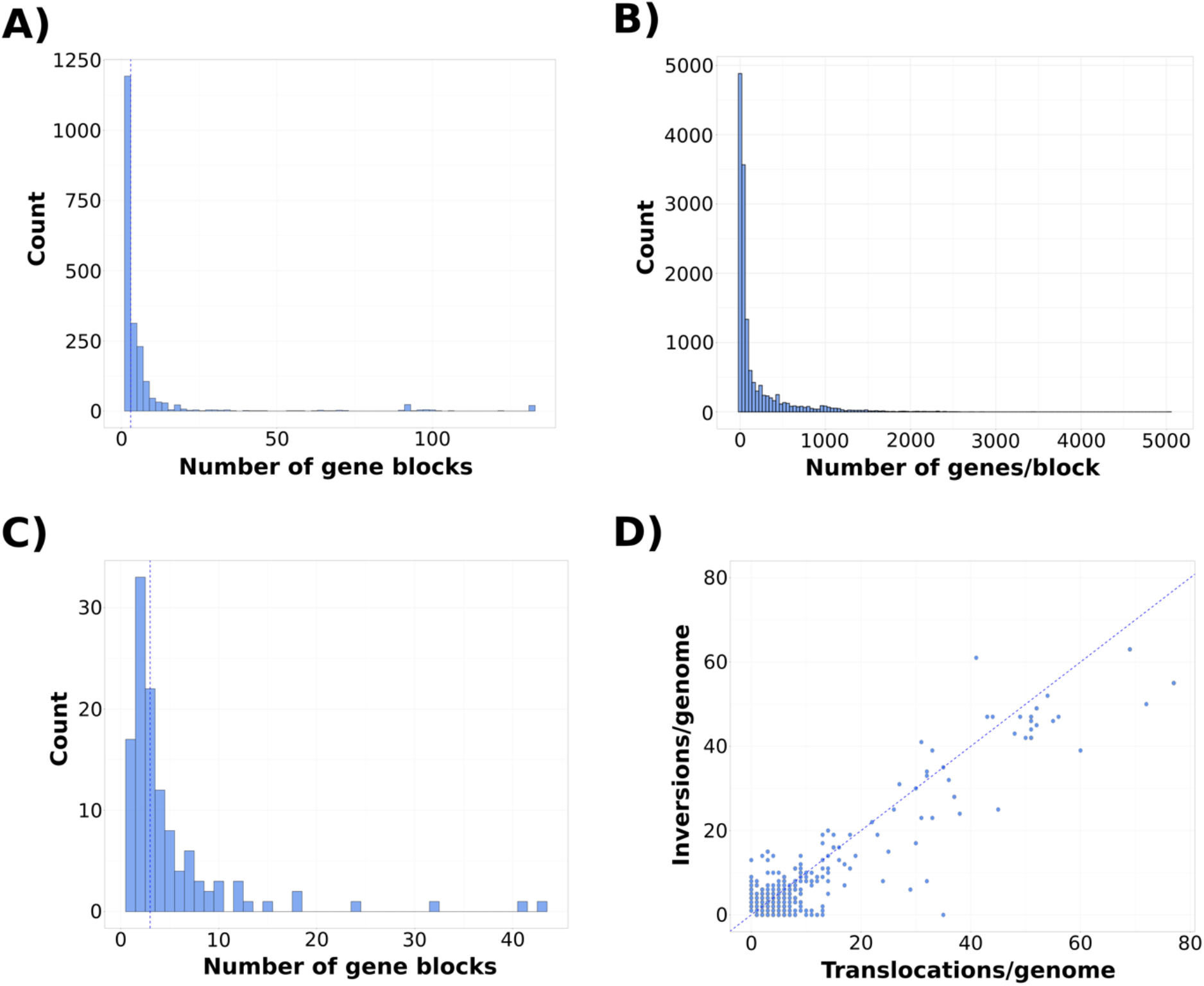
Distribution of the number of gene blocks across bacterial and archaeal species. A) Number of blocks per genome; B) Length in number of genes of rearranged blocks including accessory and core genes; C) Average number of blocks for each species; D) Number of inversions and translocations per genome.

We next investigated the relative prevalence of translocations and inversions across the rearranged genomes (Fig. 1D). Genome translocations are characterized by a shift in the order of genes without a disruption of their orientation (14, 20), whereas inversions are changes where the strand orientation of a contiguous block of genes is reversed (21). Since the net effect of inversions is a change in the orientation of genes from the leading to the lagging strand, these rearrangements are considered to be particularly deleterious (22). In agreement with this idea, our results indicate that translocations are slightly more prevalent than inversions: we found an average of four translocations vs. three inversions across our dataset (Fig. 1D).

### Location of translocations and inversions across genomes

Previous studies based on a limited set of bacterial species suggested that genome inversions may be prevalently symmetric relative to *Ori* and *Ter* (3, 21, 23). Symmetric inversions are expected to be less detrimental because they maintain the polarity of the genes in the chromosome (3, 4, 22). Moreover, DNA motifs are often polarized relative to the replication fork, and asymmetric inversions that disrupt their polarity relative to *Ori* or *Ter* can be deleterious (3, 4, 22–24). To investigate whether symmetric inversions are widespread among the species analyzed, we identified the location of the inverted gene blocks relative to *Ori* and *Ter*, whose location was determined by computing the GC skew of each genome (See Methods). For this analysis, we excluded species known for having multiple or linear chromosomes (e.g., *Vibrio* and *Burkholderia*). We further excluded a species when the identification of *Ori* and *Ter* was ambiguous based on the inspection of their GC-skew plots (see Methods).

We found that inversions are preferentially located near *Ori*, and to a lesser extent *Ter* (Fig. 2A). We next tested whether this trend could be driven by the random occurrence of inversions near *Ori* by statistically comparing the distribution of the relative position of inversions with the expectation under neutrality (See Methods). Our results show that inversions are prevalently located near *Ori* when compared with the random expectation (Wilcoxon signed rank test, *P*<0.001). Interestingly, the same trend was found in translocations (Wilcoxon signed rank test, *P*<0.001) (Fig. 2B), indicating that *Ori* is a hotspot for genomic rearrangements but not only for inversions, but also for translocations. This result suggests that rearrangements may not occur near *Ori* only because they preserve strand symmetry since this would explain the higher prevalence of inversions near *Ori*, but not the higher prevalence of translocations.

**Figure 2.**
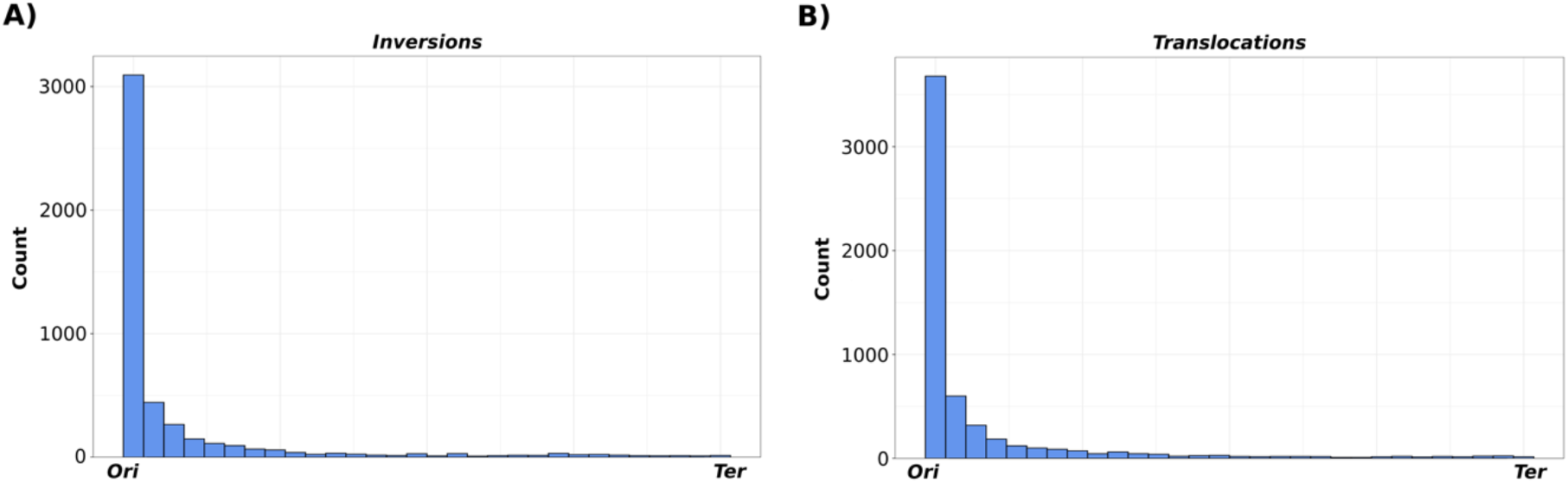
Location of genome rearrangements relative to the origin (*Ori*) and terminus (*Ter*) of replication. A) Location of the middle point of inversions relative to *Ori* and *Ter* in both chromosomal replichores; B) Location of the middle point of translocations relative to *Ori* and *Ter* in both chromosomal replichores.

### Estimation of the rate of genome rearrangement across species

Earlier studies have shown that, in general, closely related lineages show more conserved synteny than distantly related ones, and that rates of rearrangements can be estimated relative to substitution rates (20, 25). Here, we tested to what extent phylogenetic distance may be a predictor of the number of genomic rearrangements (as approximated through the number of gene blocks). We built a concatenate of core-genome alignments for each species and estimated the divergence between the pairs of genomes used to generate gene block data. In contrast to previous studies (20, 25), we observed a significant but low correlation between the evolutionary distance and the pairwise number of blocks after adjusting for core genome size (Fig. 3A; Spearman’s *Rho*=0.08; *P*<0.001). This finding indicates that rearrangements may not accumulate in genomes at a constant pace, at least over short evolutionary timescales (i.e., between strains of a species).

**Figure 3.**
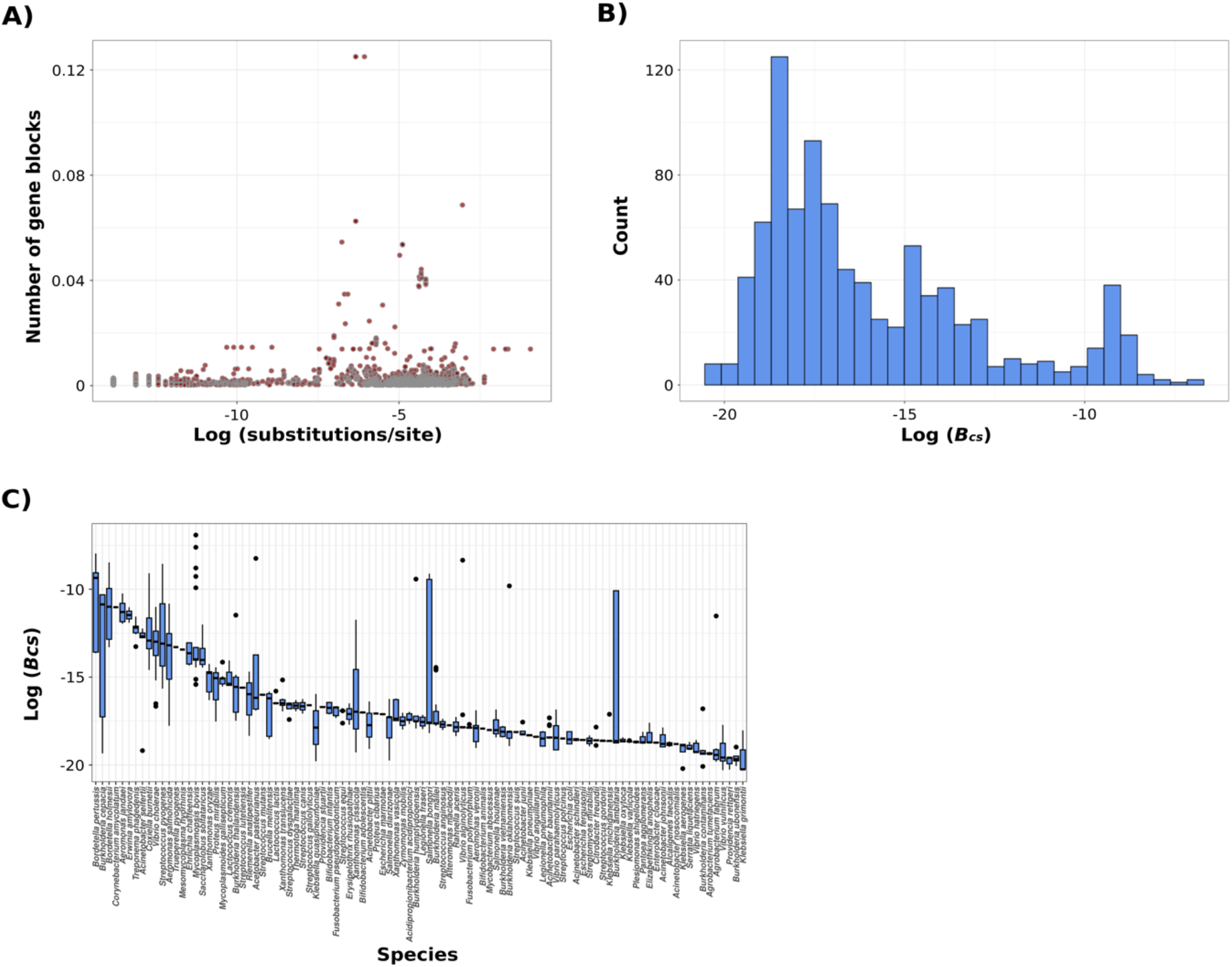
Relationship between the number of gene blocks and evolutionary distance. A) relationship between number of blocks normalized by core genome size and pairwise distance (substitutions/site) estimated from a core genome alignment for each species; B) Rearrangement rates (*B*_*sc*_) across bacterial and archaeal genomes. *B*_*sc*_ represents the number of estimated rearrangements divided by the product of the size of the core genome (number of genes), the length of the core genome alignment (in nucleotides), and the number of substitutions between genome pairs (i.e., number of blocks adjusted by the size of the core genome relative to the substitution rate). See the Methods section for more details on the estimation of *B*_*sc*_; C) Rearrangement rates (*B*_*sc*_) across species.

We further aimed at measuring at which rate genome rearrangements occur in the species of our dataset. We estimated the number of rearrangements from the number of blocks observed between genome pairs. To do so, we used simulations to establish a relationship between the number of rearrangements and the number of expected blocks resulting from these events (See Methods). Because the size of the core genome varies substantially across species, we defined the rate of rearrangement as *B*_*sc*_, which represents the number of blocks adjusted by the size of the core genome relative to the substitution rate, which allows us to compare the rate of rearrangement across species. In support of our findings of an unsteady pace of rearrangement accumulation across genomes and species, our results show a large variation of rates across genomes (Fig. 3B), but also between and within species (Fig. 3C).

### Mobile elements are likely drivers of genome rearrangements

Genome rearrangements are thought to be driven by internal processes like homologous recombination and the activity of viral recombinases, transposons, and other mobile genetic elements (MGEs) (4, 5, 26–28). To investigate the potential drivers of the genome rearrangements reported in our dataset, we analyzed the function of the core and accessory genes found in the rearranged gene blocks. We explored the prevalence of different functional categories in translocated and inverted blocks relative to the functional composition of all the genomes in our dataset (See Methods). We found a similar functional composition between translocations, inversions, non-rearranged regions, and the functional categories found across all the genomes (Fig. 4A-D). To further investigate what specific functions may have a role in genome rearrangements, we identified the COGs statistically enriched in translocations and inversions. Enriched COGs were defined as COGs presenting a Benjamini-Hochberg corrected *P*-value below 0.05 using a *χ*^2^test that compared their frequency in rearranged blocks relative to their overall expected frequency across the entire genome. Translocations and inversions showed a prevalence of enriched COGs that are involved in Carbohydrate transport and metabolism, Translation, ribosomal structure and biogenesis, Inorganic ion transport and metabolism, Signal transduction and metabolism, Coenzyme transport and metabolism, Transcription, and Energy production and conversion (adjusted *P-value*<0.05; Supplementary File 5). We further focused our analyses on the COGs that belong to the categories “Mobilome: prophages, transposons” and “Replication, recombination and repair” (adjusted *P-value*<0.05; Fig. 4) due to their expected role on genome rearrangements. The complete list of the COGs enriched in rearrangements can be found in Supplementary File 5.

**Figure 4.**
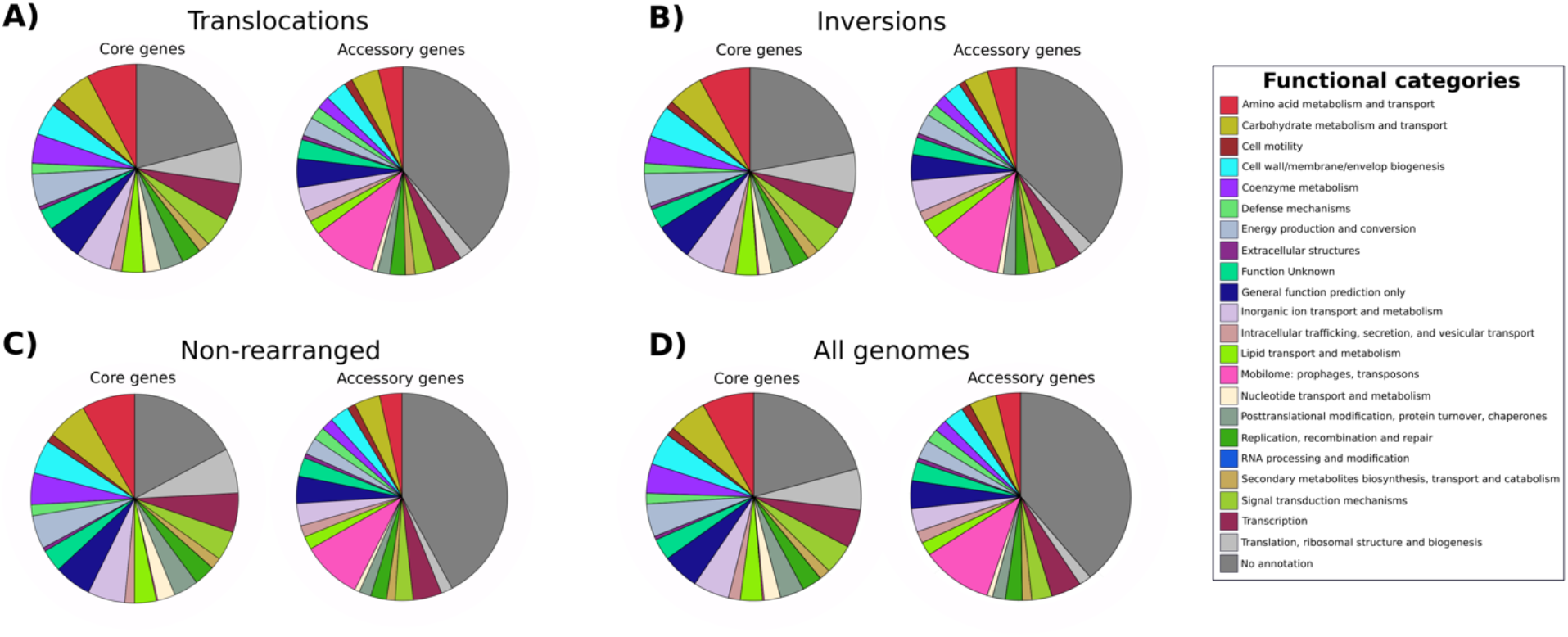
Functional annotation (COGs) of the core and accessory genes found in rearranged gene blocks relative to all the genomes.

Overall, our analysis shows that genes involved in the categories “Mobilome: prophages, transposons” and “Replication, recombination and repair” were statistically enriched in the rearranged blocks relative to the content of whole genomes (Supplementary File 5). Translocations and inversions showed a similar fraction of genes enriched in these categories. The majority of the enriched core genes in both types of rearrangements were part of the category “Replication, recombination and repair”. We found genes involved in the replication of the chromosome, such as DnaA, DnaC, and helicases. We also found an over-representation of genes encoding DNA-repair proteins like Endonuclease UvrABC ATPase subunit, Recombinational DNA repair ATPase RecF, and DNA mismatch repair protein MutH, among others. In addition, we found the enrichment of genes annotated as involved in the integration of prophages, genes involved in the duplication, recombination, and repair of DNA, and particularly genes annotated as transposases. In support of the potential role of transposons in genome rearrangements, we found a strong correlation between the total number of transposases and the number of rearranged blocks found in genomes (Spearman correlation, *Rho=*0.4; *P*<0.001, Fig. 5).

**Figure 5.**
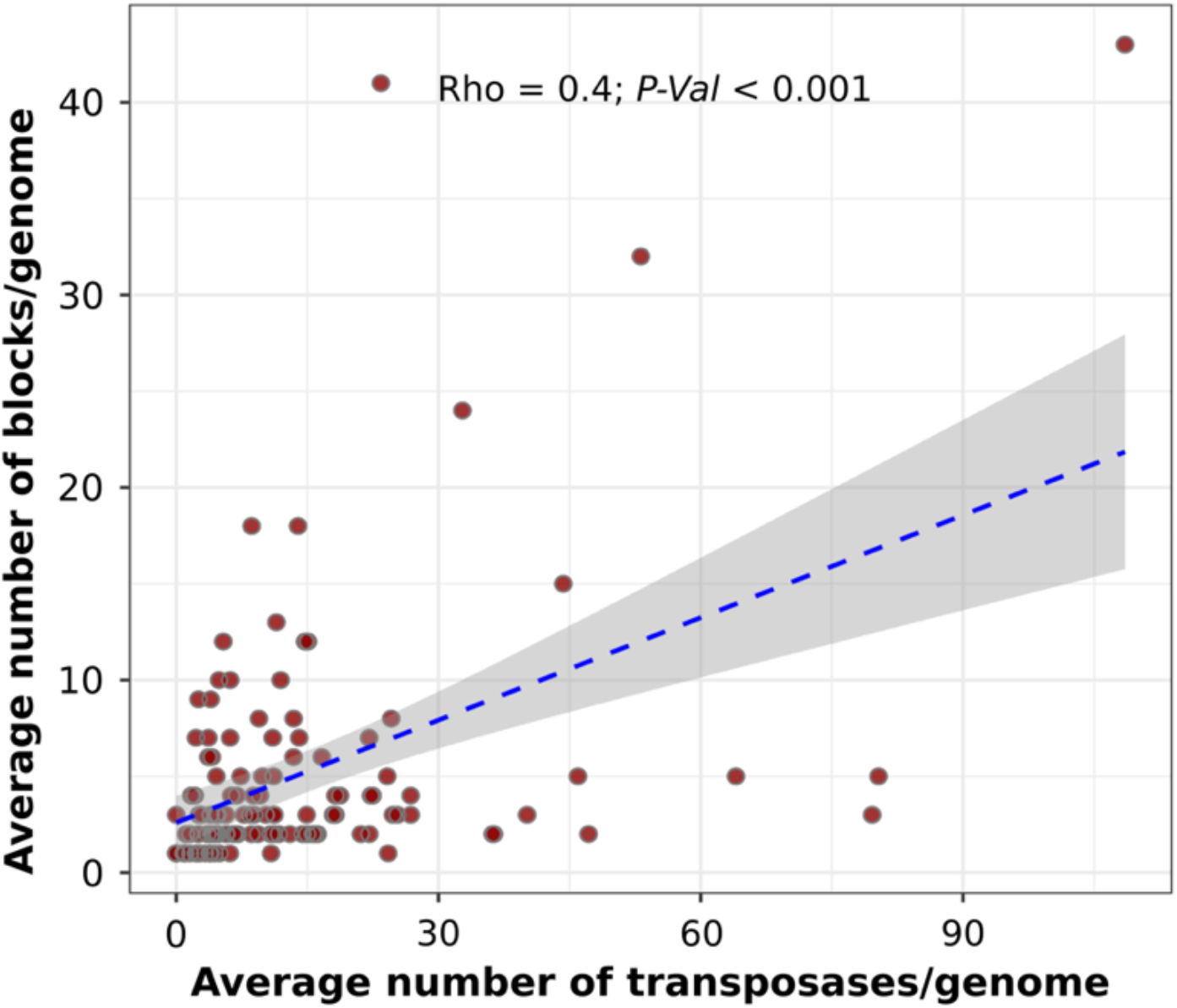
Relationship between the number of blocks found in each genome and the number of proteins annotated as transposases. Each dot represents the average number of blocks and transposases for each species. For each genome, the number of transposases was adjusted by the genome size.

The accessory genes enriched in rearranged blocks showed a large diversity of COGs in the category “Mobilome: prophages, transposons”, particularly transposases and other transposase- and phage-related genes. Examples of these genes are Transposase IS1182 family, Transposase and inactivated derivatives IS5 family, and Phage terminase large subunit, respectively. To a lesser extent, the accessory genes found in rearranged blocks were enriched in genes for the repair of DNA and recombination (Supplementary File 5).

## DISCUSSION

In this study, we explored the prevalence and the potential drivers of genomic rearrangements in a broad diversity of bacterial and archaeal species. Our results indicate that, overall, genome rearrangements are rare events within species (Fig. 1A), which is in agreement with their expected deleterious effect for cell physiology and genome stability (13, 29). On average, pairs of genomes from the same species displayed three syntenic blocks, which is typically the result of a single rearrangement (SI Appendix, Fig. S2). This finding indicates that the genomes of a given species are largely syntenic and most species display less than two rearrangement events. The rare occurrence of rearrangements in our dataset together with previous findings suggest that these events are very rare and/or that they are strongly counter selected when they occur. Similarly, we also found that when rearrangements are maintained in genomes, they span a large number of genes (Fig. 1B), suggesting that small rearrangements are as deleterious as larger ones. Note that, as opposed to some other studies (14, 16, 20, 21, 25), our analyses focused on rearrangements that occurred across short evolutionary timescales (i.e., within a species).

Although few arrangements were observed in most species, we found extreme instances at both ends of the spectrum. Some species showed a complete absence of genomic rearrangements, whereas others exhibited evidence of frequent movement of gene blocks across their genomes (Fig. 2B). Among the species where genomic rearrangements were not identified were *B. aphidicola, T. pallidum, M. morgani, B. aurantiacum, B. garinii, B. burgdorferi, and B. avium*. These species, with the exception of *B. aurantiacum*, are characterized by having symbiotic lifestyles, which likely limits the influence of external MGEs (as detailed below). Moreover, host-dependent microorganisms like *B. aphidicola* have lost genes involved in DNA repair and recombination (30–32), two processes that can drive rearrangements through intrachromosomal recombination (22). In contrast, species with a large estimated number of rearrangements like *A. salmonicida, B. pertussis, T. phagedenis, and X. oryzae*, have been reported to have a high number of transposons and other MGEs (33–37). These processes are expected to contribute to the frequent rearrangement events observed in these genomes.

Our study shows that translocations are slightly more prevalent than inversions across the species analyzed in our dataset (Fig. 1C). Although they were rare overall, inversions were found in many of the genomes where rearrangements were detected, a finding that contrasts with their expected deleterious effect on cell fitness (1). The occurrence of inversions in many of the genomes analyzed suggests that these events occasionally have a neutral or minimal effect on cellular fitness. Indeed, it was demonstrated that in *E. coli*, some inversions occurring in the same replichore are only slightly detrimental, regardless of changes in gene orientation (3, 4, 13). Interestingly, we found that not only inversions, but also translocations tend to be enriched near *Ori*, and to a lesser extent *Ter* (Fig. 2A-B). This finding contrasts with a previous study that found a higher incidence of inversions near *Ter* relative to *Ori (25)*.

Our findings indicate that *Ori* has a higher propensity for rearrangements relative to other regions of the genome. Alternatively, a more frequent relocation of genomic blocks near *Ori* may reflect an adaptive advantage to move some genes to this region through the benefit of gene dosage effect (38). DNA replication can be initiated multiple times before the first round of replication is completed, which results in multiple copies of the area proximal to *Ori* and therefore an increase in the expression of neighboring genes (39–42). Gene dosage effect may select for the relocation of genes closer to *Ori* to increase gene expression, which may constitute a rapid adaptive path to enhance gene expression. For instance, evidence suggests that changes in the location of genes from the secondary to primary chromosome in *Burkholderia*, a species known for having multiple chromosomes, can enhance the expression of genes and potentially lead to positive effects at the fitness level (43). In agreement with this idea, we observed that many of the rearranged blocks were enriched in genes involved in key cellular processes such as translation, transcription and carbohydrate metabolism. The relocation of these genes near *Ori* may confer a rapid adaptive response to some selective pressures. The rare occurrence of rearrangements at *Ter* suggests that when the relocation of genes occurs, these events are strongly counter selected in this region potentially due to the stronger deleterious effects of polarity changes or the disruption of essential gene structures. Indeed, inversions occurring near *Ter* can lead to replichores of different sizes which can then lead to asynchronous convergence of the replication forks, and cause potential issues for chromosome deconcatenation and segregation (3, 44). Moreover, *Ter* is the target for proteins that allow the correct segregation of chromosomes after replication (45), and therefore relocation in this region can negatively affect cell division. Overall, our results indicate that the translocation of genes near *Ori* may enhance cellular fitness by increasing the expression of genes, whereas inversions that do not preserve DNA polarity at *Ter* are more likely to be counter selected.

In our study, the relationship between the number of syntenic blocks and evolutionary distances was not as clear as previously reported (Fig. 3A) (42, 46). The low correlation between gene synteny and evolutionary distance is supported by the remarkable differences we observed in the rate of rearrangements across genomes and species (Fig. 3B-C). Our findings suggest that the mechanisms and evolutionary processes involved in the preservation and discontinuity of rearrangements vary across genomes and species. It must be emphasized, however, that our analyses were conducted over short evolutionary timescales and that these events are rare. It is therefore expected that our results present a much higher variance relative to rearrangement studies conducted between species and genera.

We found an enrichment of prophages and transposons in rearranged blocks, which suggests that these MGEs are key drivers of translocations and inversions in microbes (Fig. 4A-C; Figure 5; Supplementary File 5). Our findings are in close agreement with previous studies that pointed to the intrachromosomal movement of genome fragments due to the activity of phages and transposons (26, 27, 47–49). For instance, prophages in the genome of the phytopathogen *Xylella fastidiosa* can serve as anchoring points for homologous recombination and lead to major genome rearrangements (26). Another study demonstrated that recombination between homologous prophage genes can result in large chromosomal rearrangements (27). In addition, it has been shown that the proliferation of phage satellites can facilitate rearrangements in the bacterial genome (47). Similar examples have been reported for transposons (48, 50, 51). In fact, the replication cycle of some bacteriophages and other transposable elements is associated with their movement to new sites in the genome, which may cause the rearrangement of adjacent DNA sequences (49, 52).

Although we did not find significant differences in the number of blocks across microbial lifestyles (SI Appendix, Fig. S3), we observed that symbiotic bacteria that largely lack genes involved in DNA repair and recombination were among the species that showed the absence of rearrangements. These results suggest that intrinsic cellular processes such as DNA repair and recombination are additional mechanisms driving genomic rearrangements. In addition, these organisms also tend to display fewer MGEs than their free-living counterparts. The role of internal processes like DNA repair and MGEs in genomic rearrangements is in agreement with the absence of rearrangements found in endosymbionts and host-associated microbes (30, 53). Overall, our analyses suggest that genomic translocations and inversions in the microbial species that we analyzed are mostly driven by the movement and proliferation of transposable elements and prophages, and potentially processes involved in recombination and DNA repair.

## MATERIALS AND METHODS

### Genomic data collection and identification of genome rearrangements

In order to assess the prevalence of genome rearrangements in bacteria and archaea, we first compiled a genomic dataset that consisted of all the genomes available in the Genome Taxonomy Database (GTDB; release 207) (54–57) after quality filtering. Quality control consisted in retaining genomes that were 100% complete and had a contamination estimate below 5% (GTDB dataset; Supplementary File 1). We only retained genomes that were fully assembled according to assembly information available on NCBI (58) (Final dataset, Supplementary File 2). After applying these criteria, we only retained the species with more than five genomes. Our final dataset consisted of 5,238 genomes belonging to 118 bacterial species and three archaeal species.

Genomic rearrangements were detected through the identification of core gene blocks within each species. We built the core genome for each species using *CoreCruncher* (19) with the parameters -freq 90 -score 80. Gene blocks were identified using custom Python code (https://github.com/carolinaamg/prok_gene_blocks) and consisted in identifying breakpoints of consecutive core genes. Disrupted blocks were located based on pairwise comparison of the order of core genes in genomes from the same species. Blocks consisting of a single core gene were discarded as these cases could be the result of errors in orthology assignment. We defined the reference genome as the genome that led to the lowest average number of blocks across the rest of the genomes of the same species.

To assess the impact of sequencing technology on gene block estimates, the files containing the assembly statistics for each were retrieved from RefSeq. Genomes were classified as “short-read sequenced” or “long-read sequenced” based on the technology used for their sequencing process. Genomes generated with Illumina, 454, Sanger, Ion Torrent, DNBSEQ, and BGISEQ were classified as “short-read sequenced”, whereas genomes sequenced through PacBio and Nanopore were labeled as “long-read sequenced”. We only compared species that had at least five genomes sequenced with each technology (Supplementary File 2). Because our analyses showed that genomes generated through short- and long-read sequencing resulted in similar numbers of gene blocks, the genomes assembled from both sequencing technologies were retained for further analyses.

### Identification of the position of genome rearrangements relative to *Ori* and *Ter*

We identified the Origin (*Ori*) and Terminus (*Ter*) of replication in each genome by computing the cumulative GC skew (*S)* using a sliding window of 1,000 bp with *S* = (G-C)/(G+C) (59–61). Due to differences in the mutation bias of the leading and lagging strands, *Ori* and *Ter* can be identified as the global minimum and maximum of the cumulative GC skew of the DNA fragments analyzed, respectively. For this analysis, we excluded species known for having more than one chromosome or linear chromosome. We further excluded the genomes whose GC-skew plots were too ambiguous to clearly identify *Ter* and *Ori* (i.e., multiple peaks and valleys were present). Additionally, genomes with only one gene block were discarded for this analysis since they did not show any occurrence of rearrangements.

We next explored whether the location of genomic rearrangements showed a bias towards *Ori* or/and *Ter*. We identified the location of the middle of each rearrangement block relative to *Ori* and *Ter* using custom Python code. Distance values ranged from 0 to 1 and indicated the relative location from *Ori* and *Ter* for both replichores, respectively. Next, we compared each distance distribution with the expectation under neutrality (average distance of 0.5) through a Wilcoxon signed rank test. To avoid biases, we excluded genomes that had only one block (absence of genomic rearrangements) and gene blocks that appeared more than twice in our dataset.

### Estimation of genome rearrangement rates across bacterial lineages

The evolutionary distances between pairs of genomes were calculated using a nucleotide concatenated alignment consisting in the core genome of each species as reconstructed by *CoreCruncher* (see above). Alignments were generated for each core gene individually using MAFFT (62) with default parameters. Once aligned, core genes were concatenated using custom Python code (https://github.com/carolinaamg/prok_gene_blocks). We used RAxML (63) to estimate the evolutionary distance between each of the genome pairs and the corresponding number of gene blocks. Distances were computed with RAxML with the option -f x and the substitution model GTRGAMMA for all species except *Bordetella holmessi*, for which we used the model GTRGAMMAI due to the prevalence of invariable sites.

We estimated the rate of genomic rearrangements in our dataset. To generate accurate and comparable estimates of the number of rearrangements across species, we simulated genomic rearrangements and used the resulting number of blocks to fit a quadratic regression (SI Appendix, Fig.S2). We next used the equation that resulted from this regression to predict the number of rearrangements from gene block data. Then, we computed the rearrangement rate, denoted by *B*_*cs*_, as follows:

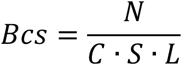

Where:

*N* is the number of predicted genomic rearrangements.

*C* is the number of genes in the core genome.

*S* is the number of nucleotide substitutions in core genome between pair genomes.

*L* is the length of core genome concatenated alignment.

In our analysis, *B*_*cs*_ is the number of blocks per substitution normalized by the core genome size and alignment length.

### Functional annotation of genomes and enrichment analysis

In order to gain insights into the potential drivers of genome rearrangements in the genomes in our dataset, we performed a functional annotation using the EggNOG database (v5.0) (64) through the hmmsearch tool available on HMMER 3.2.1 (65) with an *e*-value threshold of 10^−5^ on all proteins. Proteins with multiple annotations were filtered to keep the best-scored annotation. To find COGs enriched in genome rearrangements relative to all the genomes, we performed a *X*^2^ test on each COG found in translocations and inversions. The test matrix for each analysis consisted in the count for a given COG in rearranged blocks, the counts of other COGs in the rearranged blocks, the count of the given COG in all the genomes, and the count of other COGs in all the genomes. We excluded genomes that were composed of a single block (absence of genomic rearrangements) and blocks that were the backbone of the genome identified as the longest block.

## ACKNOWLEDGEMENTS

The authors kindly thank the members of the Bobay Lab at North Carolina State University for their thoughtful comments to improve earlier versions of the manuscript. The authors would also like to thank the Longleaf Cluster hosted at the University of North Carolina at Chapel Hill for the bioinformatic analyses performed. This study was funded by a National Institutes of Health (NIH) grant (R01GM132137) awarded to LMB and by a Simons Postdoctoral Fellowships in Marine Microbial Ecology awarded to CAMG.

## SUPPLEMENTARY FILES AND FIGURES

**Supplementary Figure 1**. Comparison of the number of blocks identified using Long-read sequenced genomes (LRSG) and Short-read sequences genomes (SRSG). Each dot represents a species.

**Supplementary Figure 2**. Quadratic relationship between the number of predicted blocks that result from simulated rearrangements.

**Supplementary Figure 3**. Number of blocks found in species across different lifestyles.

**Supplementary File 1**. Genomes retrieved from the Genome Taxonomy Database (GTDB release 207) (54–57) (GTDB dataset).

**Supplementary File 2**. Genomes used for gene block identification (Final dataset).

**Supplementary File 3**. Number of blocks identified in each genome relative to the reference genome used.

**Supplementary File 4**. Average number of blocks found in each species as well as genomes used for each species and reference genomes.

**Supplementary File 5**. Enriched and non-enriched COGs found in translocations and inversions, including accessory and core genes.

## Notes

https://github.com/carolinaamg/prok_gene_blocks

### Competing Interest Statement

The authors have declared no competing interest.

## REFERENCES

1. S. Casjens, The diverse and dynamic structure of bacterial genomes. Annu. Rev. Genet. 32, 339–377 (1998).

2. E. Darmon, D. R. F. Leach, Bacterial genome instability. Microbiol. Mol. Biol. Rev. 78, 1–39 (2014).

3. E. P. C. Rocha, The organization of the bacterial genome. Annu. Rev. Genet. 42, 211–233 (2008).

4. E. P. C. Rocha, Order and disorder in bacterial genomes. Curr. Opin. Microbiol. 7, 519–527 (2004).

5. E. P. C. Rocha, Inference and analysis of the relative stability of bacterial chromosomes. Mol. Biol. Evol. 23, 513–522 (2006).

6. D. Romero, R. Palacios, Gene amplification and genomic plasticity in prokaryotes. Annu. Rev. Genet. 31, 91–111 (1997).

7. M. B. Schmid, J. R. Roth, Gene location affects expression level in Salmonella typhimurium. J. Bacteriol. 169, 2872–2875 (1987).

8. C. Sousa, V. de Lorenzo, A. Cebolla, Modulation of gene expression through chromosomal positioning in Escherichia coli. Microbiology 143 (Pt 6), 2071–2078 (1997).

9. A. Buton, L.-M. Bobay, Evolution of Chi motifs in Proteobacteria. G3 11 (2021).

10. J. E. Rebollo, V. François, J. M. Louarn, Detection and possible role of two large nondivisible zones on the Escherichia coli chromosome. Proc. Natl. Acad. Sci. U. S. A. 85, 9391–9395 (1988).

11. S. Nolivos, et al., Co-evolution of segregation guide DNA motifs and the FtsK translocase in bacteria: identification of the atypical Lactococcus lactis KOPS motif. Nucleic Acids Res. 40, 5535–5545 (2012).

12. M. Valens, S. Penaud, M. Rossignol, F. Cornet, F. Boccard, Macrodomain organization of the Escherichia coli chromosome. EMBO J. 23, 4330–4341 (2004).

13. E. Esnault, M. Valens, O. Espéli, F. Boccard, Chromosome structuring limits genome plasticity in Escherichia coli. PLoS Genet. 3, e226 (2007).

14. E. R. Tillier, R. A. Collins, Genome rearrangement by replication-directed translocation. Nat. Genet. 26, 195–197 (2000).

15. A. U. Kresse, S. D. Dinesh, K. Larbig, U. Römling, Impact of large chromosomal inversions on the adaptation and evolution of Pseudomonas aeruginosa chronically colonizing cystic fibrosis lungs. Mol. Microbiol. 47, 145–158 (2003).

16. C. N. Merrikh, H. Merrikh, Gene inversion potentiates bacterial evolvability and virulence. Nat. Commun. 9, 4662 (2018).

17. N. Campo, M. J. Dias, M.-L. Daveran-Mingot, P. Ritzenthaler, P. Le Bourgeois, Chromosomal constraints in Gram-positive bacteria revealed by artificial inversions. Mol. Microbiol. 51, 511–522 (2004).

18. P.-A. Chaumeil, A. J. Mussig, P. Hugenholtz, D. H. Parks, GTDB-Tk: a toolkit to classify genomes with the Genome Taxonomy Database. Bioinformatics 36, 1925–1927 (2019).

19. C. D. Harris, E. L. Torrance, K. Raymann, L.-M. Bobay, CoreCruncher: Fast and Robust Construction of Core Genomes in Large Prokaryotic Data Sets. Mol. Biol. Evol. 38, 727–734 (2021).

20. E. Belda, A. Moya, F. J. Silva, Genome rearrangement distances and gene order phylogeny in gamma-Proteobacteria. Mol. Biol. Evol. 22, 1456–1467 (2005).

21. J. A. Eisen, J. F. Heidelberg, O. White, S. L. Salzberg, Evidence for symmetric chromosomal inversions around the replication origin in bacteria. Genome Biol. 1, RESEARCH0011 (2000).

22. A. E. Darling, I. Miklós, M. A. Ragan, Dynamics of genome rearrangement in bacterial populations. PLoS Genet. 4, e1000128 (2008).

23. P. Mackiewicz, D. Mackiewicz, M. Kowalczuk, S. Cebrat, Flip-flop around the origin and terminus of replication in prokaryotic genomes. Genome Biol. 2, INTERACTIONS1004 (2001).

24. C. W. Hill, J. A. Gray, Effects of chromosomal inversion on cell fitness in Escherichia coli K-12. Genetics 119, 771–778 (1988).

25. M. Suyama, P. Bork, Evolution of prokaryotic gene order: genome rearrangements in closely related species. Trends Genet. 17, 10–13 (2001).

26. H. Brüssow, C. Canchaya, W.-D. Hardt, Phages and the evolution of bacterial pathogens: from genomic rearrangements to lysogenic conversion. Microbiol. Mol. Biol. Rev. 68, 560–602, table of contents (2004).

27. I. Nakagawa, et al., Genome sequence of an M3 strain of Streptococcus pyogenes reveals a large-scale genomic rearrangement in invasive strains and new insights into phage evolution. Genome Res. 13, 1042–1055 (2003).

28. A. Toussaint, P. A. Rice, Transposable phages, DNA reorganization and transfer. Curr. Opin. Microbiol. 38, 88–94 (2017).

29. H. Hendrickson, J. G. Lawrence, Selection for chromosome architecture in bacteria. J. Mol. Evol. 62, 615–629 (2006).

30. I. Tamas, et al., 50 million years of genomic stasis in endosymbiotic bacteria. Science 296, 2376–2379 (2002).

31. R. C. H. J. van Ham, et al., Reductive genome evolution in Buchnera aphidicola. Proc. Natl. Acad. Sci. U. S. A. 100, 581–586 (2003).

32. N. A. Moran, A. Mira, The process of genome shrinkage in the obligate symbiont Buchnera aphidicola. Genome Biol. 2, RESEARCH0054 (2001).

33. M. Mushtaq, S. Manzoor, M. Pringle, A. Rosander, E. Bongcam-Rudloff, Draft genome sequence of “Treponema phagedenis” strain V1, isolated from bovine digital dermatitis. Stand. Genomic Sci. 10, 67 (2015).

34. A. J. King, T. van Gorkom, H. G. J. van der Heide, A. Advani, S. van der Lee, Changes in the genomic content of circulating Bordetella pertussis strains isolated from the Netherlands, Sweden, Japan and Australia: adaptive evolution or drift? BMC Genomics 11, 64 (2010).

35. K. L. Sealey, T. Belcher, A. Preston, Bordetella pertussis epidemiology and evolution in the light of pertussis resurgence. Infect. Genet. Evol. 40, 136–143 (2016).

36. S. J. Charette, Microbe profile: an opportunistic pathogen with multiple personalities. Microbiology 167 (2021).

37. S. L. Salzberg, et al., Genome sequence and rapid evolution of the rice pathogen Xanthomonas oryzae pv. oryzae PXO99A. BMC Genomics 9, 204 (2008).

38. E. Couturier, E. P. C. Rocha, Replication-associated gene dosage effects shape the genomes of fast-growing bacteria but only for transcription and translation genes. Mol. Microbiol. 59, 1506–1518 (2006).

39. Timing of initiation of chromosome replication in individual Escherichia coli cells. EMBO J. 5, 3074 (1986).

40. H. J. Nielsen, B. Youngren, F. G. Hansen, S. Austin, Dynamics of Escherichia coli chromosome segregation during multifork replication. J. Bacteriol. 189, 8660–8666 (2007).

41. O. O. Bochkareva, E. V. Moroz, I. I. Davydov, M. S. Gelfand, Genome rearrangements and selection in multi-chromosome bacteria Burkholderia spp. BMC Genomics 19, 965 (2018).

42. M. Noureen, I. Tada, T. Kawashima, M. Arita, Rearrangement analysis of multiple bacterial genomes. BMC Bioinformatics 20, 631 (2019).

43. J. D. Morrow, V. S. Cooper, Evolutionary effects of translocations in bacterial genomes. Genome Biol. Evol. 4, 1256–1262 (2012).

44. H. Niki, S. Hiraga, Polar localization of the replication origin and terminus in Escherichia coli nucleoids during chromosome partitioning. Genes Dev. 12, 1036–1045 (1998).

45. E. Crozat, et al., Post-replicative pairing of sister ter regions in Escherichia coli involves multiple activities of MatP. Nat. Commun. 11, 3796 (2020).

46. M. B. Pietro Liò Vincent Lacroix and Marie-France Sagot, Short and long-term genome stability analysis of prokaryotic genomes. BMC Genomics 14, 1–18 (2013).

47. J. A. Moura de Sousa, E. P. C. Rocha, To catch a hijacker: abundance, evolution and genetic diversity of P4-like bacteriophage satellites. Philos. Trans. R. Soc. Lond. B Biol. Sci. 377, 20200475 (2022).

48. J. Parkhill, et al., Comparative analysis of the genome sequences of Bordetella pertussis, Bordetella parapertussis and Bordetella bronchiseptica. Nat. Genet. 35, 32–40 (2003).

49. P. Nevers, H. Saedler, Transposable genetic elements as agents of gene instability and chromosomal rearrangements. Nature 268, 109–115 (1977).

50. S. Wu, P. Tian, T. Tan, Genomic landscapes of bacterial transposons and their applications in strain improvement. Appl. Microbiol. Biotechnol. 106, 6383–6396 (2022).

51. J. Vandecraen, M. Chandler, A. Aertsen, R. Van Houdt, The impact of insertion sequences on bacterial genome plasticity and adaptability. Crit. Rev. Microbiol. 43, 709–730 (2017).

52. J. A. Shapiro, Molecular model for the transposition and replication of bacteriophage Mu and other transposable elements. Proc. Natl. Acad. Sci. U. S. A. 76, 1933–1937 (1979).

53. M. Naito, T. E. Pawlowska, The role of mobile genetic elements in evolutionary longevity of heritable endobacteria. Mob. Genet. Elements 6, e1136375 (2016).

54. D. H. Parks, et al., GTDB: an ongoing census of bacterial and archaeal diversity through a phylogenetically consistent, rank normalized and complete genome-based taxonomy. Nucleic Acids Res. 50, D785–D794 (2022).

55. C. Rinke, et al., A standardized archaeal taxonomy for the Genome Taxonomy Database. Nat Microbiol 6, 946–959 (2021).

56. D. H. Parks, et al., A complete domain-to-species taxonomy for Bacteria and Archaea. Nat. Biotechnol. 38, 1079–1086 (2020).

57. D. H. Parks, et al., A standardized bacterial taxonomy based on genome phylogeny substantially revises the tree of life. Nat. Biotechnol. 36, 996–1004 (2018).

58. E. W. Sayers, et al., Database resources of the national center for biotechnology information. Nucleic Acids Res. 50, D20–D26 (2022).

59. A. Grigoriev, Analyzing genomes with cumulative skew diagrams. Nucleic Acids Res. 26, 2286–2290 (1998).

60. J. Lu, S. L. Salzberg, SkewIT: The Skew Index Test for large-scale GC Skew analysis of bacterial genomes. PLoS Comput. Biol. 16, e1008439 (2020).

61. J. Bohlin, et al., Analysis of intra-genomic GC content homogeneity within prokaryotes. BMC Genomics 11, 464 (2010).

62. K. Katoh, D. M. Standley, MAFFT multiple sequence alignment software version 7: improvements in performance and usability. Mol. Biol. Evol. 30, 772–780 (2013).

63. A. Stamatakis, RAxML version 8: a tool for phylogenetic analysis and post-analysis of large phylogenies. Bioinformatics 30, 1312–1313 (2014).

64. J. Huerta-Cepas, et al., eggNOG 5.0: a hierarchical, functionally and phylogenetically annotated orthology resource based on 5090 organisms and 2502 viruses. Nucleic Acids Res. 47, D309–D314 (2019).

65. S. R. Eddy, Accelerated Profile HMM Searches. PLoS Comput. Biol. 7, e1002195 (2011).

